# Hyperlipidemia abolishes, but immune balancing by DNase-I restores neuroprotection by MSC-derived extracellular vesicles

**DOI:** 10.64898/2026.08.04.742906

**Authors:** Chen Wang, Tobias Tertel, Yiqiao Zhang, Yanis Mouloud, Xiaolong Liu, Nina Hagemann, Ayan Mohamud Yusuf, Aurel Popa-Wagner, Matthias Gunzer, Bernd Giebel, Dirk M. Hermann

**Affiliations:** Department of Neurology, University of Duisburg-Essen, Essen, Germany; Center for Translational and Behavioral Neurosciences, University of Duisburg-Essen, Essen, Germany; Institute of Transfusion Medicine, University Hospital Essen, University of Duisburg-Essen, Essen, Germany; Center of Normal and Pathological Ageing, University of Medicine and Pharmacy Craiova, Craiova, Romania; Institute of Experimental Immunology and Imaging, University Hospital Essen, University of Duisburg-Essen, Essen, Germany

**Author notes:** <u>Correspondence:</u> Prof. Dirk M. Hermann, MD, Department of Neurology, University Hospital Essen, University of Duisburg-Essen, Hufelandstr. 55, 45147 Essen, Germany.

**Keywords:** Ischemic stroke, hypercholesterolemia, hyperlipidemia, innate immunity, neutrophil, monocyte/ macrophage, DNase-I

## Abstract

**Background:** Owing to their potent immunomodulatory properties, mesenchymal stromal cell (MSC)-derived small extracellular vesicles (EVs) have emerged as promising neuroprotective treatments for ischemic stroke. Preclinical studies using MSC-EVs have mainly been performed in young, otherwise healthy rodents. Stroke patients frequently carry vascular risk factors and comorbidities. We herein investigated whether MSC-EVs retain neuroprotective activity in hyperlipidemic mice on cholesterol-rich Western diet.

**Methods:** Male C57BL/6J mice were exposed to regular normal diet or Western diet for 6 weeks. At the age of 9-10 weeks, mice were exposed to transient intraluminal middle cerebral artery occlusion (MCAO). Vehicle or MSC-EVs (2x10^6^ or 6x10^6^ cell equivalents) were intravenously administered immediately after reperfusion, and vehicle or rosuvastatin (5 mg/kg/day) were intraperitoneally applied starting immediately after or seven days before MCAO. Neurological deficits, ischemic injury, and immune responses were evaluated up to 72 hours post-ischemia. To investigate the hyperlipidemia-associated immune dysregulation, mice received DNase-I before or immediately after MCAO. In defined subgroups, monocytes/ macrophages or neutrophils were additionally depleted by clodronate liposomes or anti-Ly6G antibodies, respectively.

**Results:** In contrast to normolipidemic control mice, MSC-EVs failed to induce post-ischemic neuroprotection in hyperlipidemic mice. Neither MSC-EV dose escalation nor rosuvastatin co-treatment restored the therapeutic efficacy of MSC-EVs. Hyperlipidemia induced systemic innate immune dysregulation characterized by reduced monocyte/ macrophage activation, increased neutrophil activation, and elevated circulating cell-free DNA. DNase-I treatment before, but not after MCAO reversed these immune abnormalities and restored neuroprotection by MSC-EVs, decreasing neurological deficits, infarct volume and brain edema. Depletion of either monocytes/ macrophages or neutrophils abolished the neuroprotective effects of MSC-EVs in DNase-I-pretreated hyperlipidemic mice.

**Conclusions:** Immune dysregulation abolishes MSC-EV-induced neuroprotection after ischemic stroke in hyperlipidemic mice. DNase-I priming restores MSC-EV responsiveness through mechanisms critically involving monocyte/ macrophage and neutrophil rebalancing. Our data highlight the host immune status as determinant of EV therapeutic efficacy.

## Introduction

Recent advances in reperfusion therapies (that is, thrombolysis and endovascular thrombectomy ^1,2^) have reinvigorated efforts in the development of neuroprotective therapies. The reperfused brain is prone to neuroinflammatory responses associated with brain leukocyte infiltrates, which exacerbate ischemic damage.^3,4^ Mesenchymal stromal cell (MSC)-derived extracellular vesicles (EVs) have emerged as potent antiinflammatory treatments with advantages over pharmacological and cell-based therapies, owing to their versatile actions, simple handling and high safety profile.^5^ We and others have demonstrated that MSC-EVs improve neurological recovery, reduce infarct volume, increase periinfarct angiogenesis and increase neuronal survival and plasticity in young and aged mice and rats.^6-9^ Mechanistically, our previous studies identified peripheral monocytes/ macrophages and neutrophils as indispensable mediators of MSC-EV-induced neuroprotection, highlighting the importance of systemic immune responses as critical factors influencing MSC-EV efficacy.^7,10,11^

Hyperlipidemia, one of the most prevalent vascular risk factors found in about 50% of ischemic stroke patients,^12,13^ profoundly alters systemic immunity and worsens stroke outcome.^13,14^ Mice fed with a cholesterol-rich Western diet exhibit monocyte/ macrophage dysfunction, excessive neutrophil activation, and increased circulating cell-free DNA levels in the blood.^15-17^ When exposed to ischemic stroke by transient middle cerebral artery occlusion (MCAO), neurological impairments, infarct volume, brain edema, blood-brain barrier breakdown, and brain leukocyte (most notably neutrophil) infiltrates were increased in hyperlipidemic mice compared with normolipidemic mice.^13,14,18-20^ Considering the crucial role of monocytes/ macrophages and neutrophils in mediating neuroprotective MSC-EV actions,^7,10,11^ we asked if the dysregulation of immune balance might compromise their neuroprotective properties. Indeed, previous studies found that the recovery-promoting effects of other restorative therapies, i.e., vascular endothelial growth factor (VEGF) or neural precursor cells (NPCs), were impaired in hyperlipidemic compared with normolipidemic MCAO mice.^18,21^ To explore the effects of MSC-EVs under conditions of hyperlipidemia, we fed mice with a cholesterol-rich Western diet for 6 weeks and then exposed them to transient MCAO. Following the observation that MSC-EVs failed to induce neuroprotection in hyperlipidemic MCAO mice, we evaluated underlying mechanisms and explored rescue strategies to restore MSC-EV responses.

## Materials and Methods

### Legal issues, randomization and statistical planning

A detailed description of all methods is provided in the Supplemental Material. Experiments were performed with local government approval (State Office for Consumer Protection and Food North Rhine-Westphalia, Recklinghausen) in accordance to EU (Directive 2010/63/EU) and local institutional guidelines for the care and use of laboratory animals in accordance with ARRIVE guidelines for reporting animal experiments.^22^ Experiments were strictly randomized and blinded. Statistical planning assumed an alpha error of 5% and a beta error (1–statistical power) of 20%.

### MSC culture, EV preparation, and characterization

The MSC-EV preparations used in the present study were identical to our previous studies.^7,8,11,23,24^ Briefly, small EVs were isolated (1) from conditioned media of clonally expanded immortalized MSCs (ciMSCs) cultured under normoxic conditions (21% O_2_; MSC-EV preparation I), (2) from conditioned media of primary MSCs cultured under normoxic conditions (MSC-EV preparation II) or (3) from conditioned media of primary MSCs cultured under hypoxic conditions (1% O_2_; MSC-EV preparation III) using an optimized polyethylene glycol 6000 precipitation protocol followed by ultracentrifugation, as previously reported.^7^ These preparations had previously been shown to be neuroprotective in normolipidemic mice exposed to MCAO.^7,8,11^ MSC-EV preparations were characterized in accordance with the Minimal Information for Studies of Extracellular Vesicles 2018 (MISEV2018) guidelines,^25^ including nanoparticle tracking analysis (NTA), bicinchoninic acid (BCA) assay, and imaging flow cytometry for EV surface markers.

### Quantification of plasma double-stranded DNA levels

Circulating cell-free double-stranded DNA (dsDNA) levels were quantified using the Qubit dsDNA HS Assay Kit (Q32851; Thermo Fisher Scientific) according to the manufacturer’s instructions and analyzed using a Qubit 4 Fluorometer (Thermo Fisher Scientific).

### Focal cerebral ischemia

Male C57BL/6J mice (3-4 weeks old; Harlan Laboratories, Darmstadt, Germany) were kept on either standard normal (V1534-300; ssniff Spezialdiäten, Soest, Germany; 3.3% fat) or Western diet (E15721-34; ssniff Spezialdiäten; 21.1% fat) for 6 weeks.^21^ Plasma levels of total cholesterol, low-density lipoprotein (LDL), high-density lipoprotein (HDL), triglycerides, and glucose were measured by the Central Laboratory, Research and Teaching Unit, of the University Hospital Essen using enzymatic assays. Focal cerebral ischemia was induced by transient intraluminal MCAO as previously described.^7,8^ Briefly, mice were anesthetized with 1.5% isoflurane (30% O_2_, remainder N_2_O). Rectal temperature was maintained between 36.5 and 37.0 °C using a feedback-controlled heating system (Fluovac, Harvard apparatus, Holliston, MA, U.S.A.). Cerebral blood flow was recorded by laser Doppler flowmetry (Perimed, Stockholm, Sweden) above the core of the middle cerebral artery territory. A silicon-coated 7.0 nylon monofilament (Doccol Corporation, Sharon, MA, U.S.A.) was introduced through a small incision into the common carotid artery and advanced to the carotid bifurcation for MCAO. After 30 min, reperfusion was initiated by monofilament removal.

### Administration of MSC-EVs, rosuvastatin, and DNase-I

Immediately after reperfusion onset, mice received an intravenous injection of either vehicle (200 μl normal saline) or MSC-EVs (2x10^6^ or 6x10^6^ cell equivalents in 200 μl normal saline) via the tail vein.^7,8,11^ Rosuvastatin (5 mg/kg; dissolved in 200 μl normal saline; SML1264; Sigma-Aldrich, Taufkirchen, Germany) was administered intraperitoneally 0, 24, and 48 hours after reperfusion or once daily for 7 consecutive days before MCAO, as previously reported.^26^ For DNase-I preconditioning, mice received an intravenous injection of 100 µl vehicle (normal saline) or DNase-I (10 μg; 11284932001; Roche Diagnostics, Mannheim, Germany) one day before MCAO, followed by two intraperitoneal injections of 200 µl vehicle or DNase-I (50 μg) administered at 12-hour intervals.^27^ For post-ischemic DNase-I treatment, 100 µl vehicle or DNase-I (10 μg) was intravenously applied immediately after reperfusion shortly before MSC-EV delivery.^27^

### Depletion of monocytes/ macrophages and neutrophils

Peripheral monocytes/ macrophages were depleted using clodronate liposomes (Liposoma, Amsterdam, Netherlands), administered intravenously 24 hours before (50 mg/kg) and 24 and 48 hours after (30 mg/kg) MCAO, as previously described.^11^ We have confirmed in our recent study that clodronate liposomes effectively deplete circulating monocytes/ macrophages without affecting other phagocytic cell populations, including neutrophils and dendritic cells.^11^ Neutrophils were depleted by intraperitoneal administration of 200 μg anti-Ly6G antibody (1A8; BE0075-1; BioXCell, Lebanon, NH, U.S.A.) 24 hours before and 24 hours after MCAO.^7,11^

### Analysis of neurological deficits, infarct volume, and brain edema

Neurological deficits were evaluated at 24, 48, and 72 hours after MCAO using the Clark score, as previously described.^7,8^ 20-μm-thick coronal brain cryostat sections collected at 1 mm intervals across the forebrain were stained with cresyl violet. In all sections, infarct area was determined using the indirect method that corrects for brain swelling by subtracting area of healthy tissue of the ischemic hemisphere from that of the contralesional hemisphere using Image J software (National Institute of Health, Bethesda, MD, U.S.A.). Infarct volume was determined by integrating infarct areas from all brain levels, and brain edema was measured as ratio of ipsilateral to contralateral hemisphere volume.

### Immunohistochemistry

20-μm-thick coronal brain sections obtained from the rostrocaudal level of the bregma (i.e., the core of the middle cerebral artery territory) were immunolabeled for NeuN (neuronal marker), CD45 (leukocyte marker), Ly6G (neutrophil marker), CD31 (vascular endothelium marker), ICAM-1 (intercellular adhesion molecule-1, an inflammation marker on endothelial cells), collagen-IV (a marker of ischemic microvessels), GPIbα (glycoprotein Ibα, a platelet marker), or extravasated serum IgG (a blood-brain barrier permeability marker). Nuclei were counterstained with Hoechst 33342. NeuN stainings were costained with terminal deoxynucleotidyl transferase dUTP nick end labeling (TUNEL) (12156792910; Roche Diagnostics, Mannheim, Germany). Injured NeuN^+^/ TUNEL^+^ neurons, ICAM-1^+^ microvessels, CD45^+^ leukocytes, Ly6G^+^ neutrophils, GPIbα^+^ microvascular thrombi, and IgG extravasation were evaluated in the ischemic striatum by counting cell numbers, determining cell densities, or measuring optical densities.

### Flow cytometry of leukocytes

Single cell suspensions for flow cytometry analysis were prepared as previously described.^7,8^ Cell suspensions were stained with antibody cocktails listed in Supplemental Table S2 and analyzed by a CytoFLEX flow cytometer (Beckman-Coulter) using Kaluza software V2.2 (Beckman-Coulter).

### Statistical analysis

Statistical analysis was performed using GraphPad Prism (version 10.4.1; GraphPad Software, San Diego, California U.S.A.). Normal distribution was assessed in all datasets using Shapiro-Wilk normality tests. Normally distributed data were analyzed by one-way ANOVA followed by LSD post hoc tests, two-way ANOVA followed by LSD post hoc tests (comparisons between ≥ 3 groups) or unpaired or paired two-tailed Student’s t tests (comparisons between 2 groups), as adequate. Non-normally distributed data were evaluated by Kruskal-Wallis tests followed by Dunn’s multiple comparison tests (comparisons between ≥ 3 groups) or two-tailed Mann-Whitney U tests (comparisons between 2 groups). Data were presented as box plots with median ± interquartile ranges with minimum and maximum values as whiskers and individual data points as dots. P<0.05 was considered statistically significant.

## Results

### MSC-EVs fail to induce neuroprotection in hyperlipidemic mice on Western diet

Consistent with our previous studies,^13,21^ plasma levels of total cholesterol, low-density lipoprotein (LDL), high-density lipoprotein (HDL), and glucose were significantly increased in hyperlipidemic mice on Western diet compared to normolipidemic mice on normal diet (Figure 1A). We thus exposed normolipidemic and hyperlipidemic mice to MCAO and administered small EVs (2x10^6^ cell equivalents) isolated from conditioned media of clonally expanded immortalized MSCs (ciMSCs) cultured under normoxic (21% O_2_; MSC-preparation I) condition or from conditioned media of primary MSCs cultured under normoxic (MSC-preparation II) or hypoxic (1% O_2_; MSC preparation III) conditions immediately post-MCAO (Figure 1B). MSC-EV preparations induced ischemic neuroprotection in normolipidemic mice, but failed to do so in hyperlipidemic mice (Figure 1C, D). This loss of neuroprotection was noted for EVs isolated from ciMSCs cultured under normoxic conditions (MSC-EV preparation I), EVs isolated from primary MSCs cultured under normoxic conditions (MSC-EV preparation II) and EVs isolated from primary MSCs cultured under hypoxic conditions (MSC-EV preparation III) (Figure 1C, D). As such, all three MSC-EV preparations reproducibly reduced neurological deficits, assessed using the Clark score, and infarct volume, assessed by cresyl violet staining, in normolipidemic mice, but not in hyperlipidemic mice post-MCAO.

**Figure 1.**
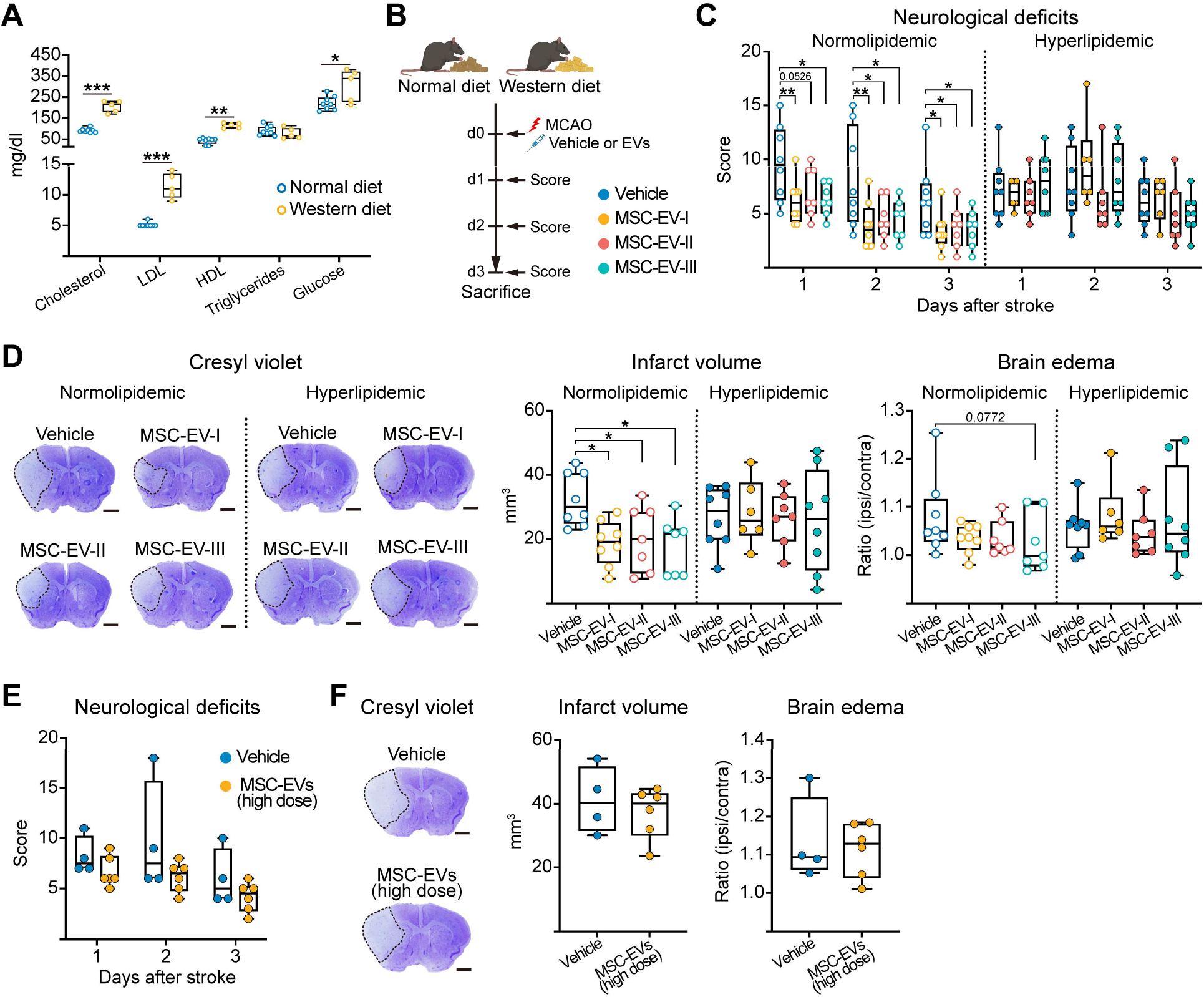
Mesenchymal stromal cell (MSC)-derived extracellular vesicles (EVs) fail to induce ischemic neuroprotection in hyperlipidemic mice on Western diet. (**A**) Plasma levels of total cholesterol, low-density lipoprotein (LDL), high-density lipoprotein (HDL), triglycerides, and glucose in non-ischemic mice after 6 weeks of standard normal diet (ND) or cholesterol-rich Western diet (WD) feeding. (**B**) Experimental design: Mice were fed with the two diets for 6 weeks and subjected to MCAO. Vehicle or MSC-EVs (2x10^6^ cell equivalents) were intravenously administered immediately after reperfusion onset. Neurological deficits were evaluated at 24, 48, and 72 hours using the Clark score, followed by animal sacrifice at 72 hours post-MCAO (created with https://BioRender.com). (**C**) Neurological deficits using the Clark score assessed at 24, 48, and 72 hours after MCAO and (**D**) infarct volume and hemisphere brain edema evaluated by cresyl violet staining at 72 hours post-MCAO in mice treated with vehicle or three different MSC-EV preparations (MSC-EV-I: isolated from clonally expanded immortalized MSCs (ciMSCs) cultured under normoxia (21% O_2_); MSC-EV-II: isolated from primary MSCs cultured under normoxia; MSC-EV-III: isolated from primary MSCs cultured under hypoxia (1% O_2_)). (**E**) Neurological deficits, (**F**) infarct volume and brain edema evaluated in MCAO mice receiving vehicle or high-dose MSC-EVs (6x10^6^ cell equivalents; preparation MSC-EV-I). Representative cresyl violet stainings are shown (in (**D, F**)). Data were evaluated by unpaired t tests or Mann-Whitney U tests (in (**A, E, F**)) or one-way ANOVA with LSD post hoc tests (in (**C, D**)). *p<0.05, **p<0.01, ***p<0.001 (n=5-8 mice/group (in (**A**)), n=6-8 mice/group (in (**C, D**)), n=4-6 mice/group (in (**E, F**)). Scale bars: 1 mm (in (**D, F**)).

### EV dose escalation and HMG-CoA reductase inhibition fail to rescue MSC-EV-induced neuroprotection in hyperlipidemic mice

We next wondered whether elevated lipoprotein levels masked the neuroprotective effects of MSC-EVs, e.g., by impairing EV binding and uptake by recipient cells.^28,29^ We therefore explored whether dose escalation could restore the neuroprotective effects of MSC-EVs. Importantly, a 3-fold higher dose of MSC-EVs (6x10^6^ cell equivalents) failed to salvage functional or structural neuroprotection by MSC-EVs in hyperlipidemic MCAO mice (Figure 1E, F). Since hyperlipidemic patients commonly receive cholesterol-lowering HMG-CoA reductase inhibitors (i.e., statins), we next examined whether add-on treatment with rosuvastatin (5 mg/kg/day; i.p.) could reestablish MSC-EV-induced neuroprotection. Of note, rosuvastatin again did not restore neuroprotection, neither when administered immediately after MCAO (Figure 2A-C) nor over 7 days before MCAO (Figure 2D-F).

**Figure 2.**
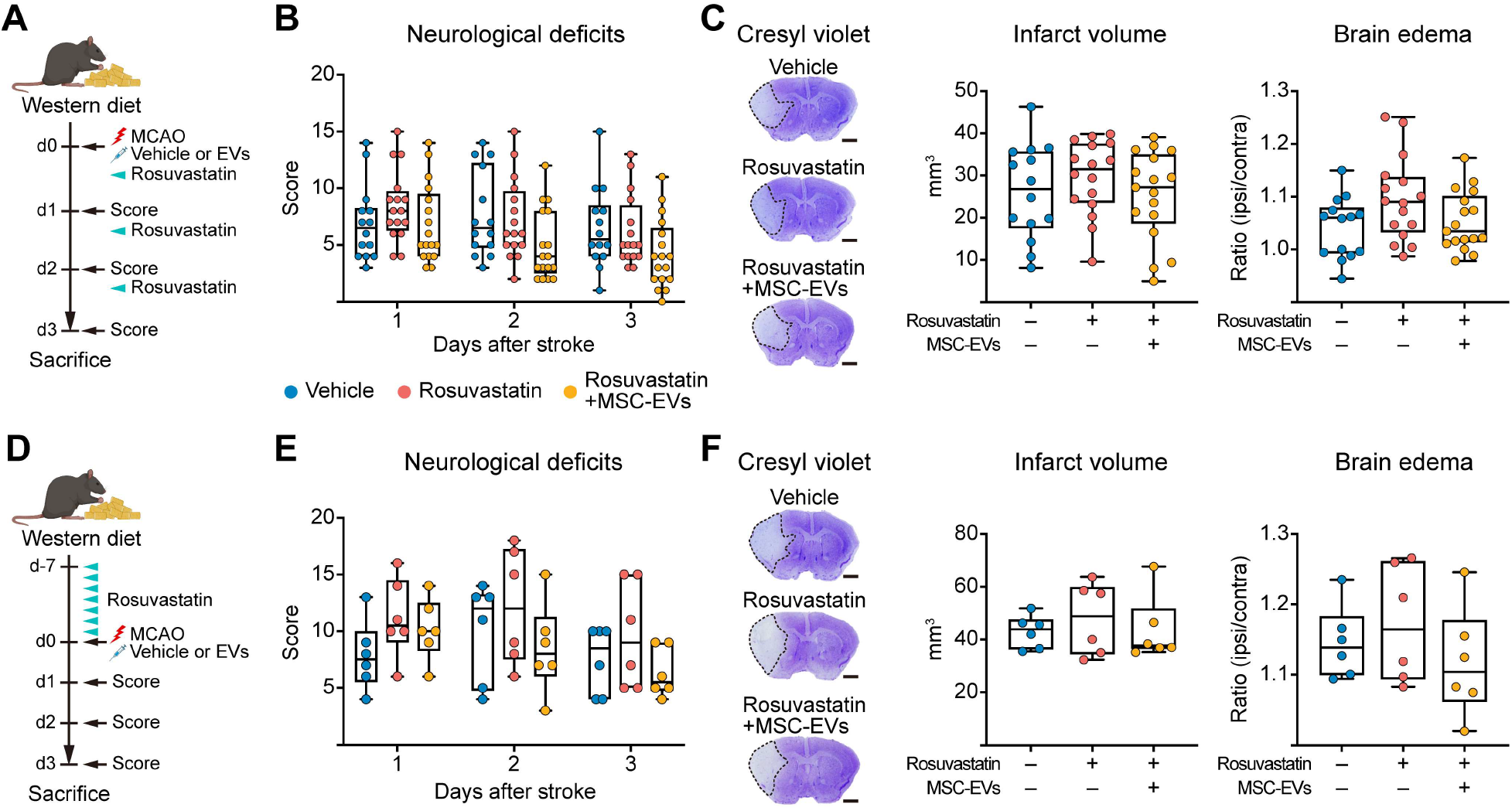
Add-on treatment with rosuvastatin fails to rescue MSC-EV-induced ischemic neuroprotection in hyperlipidemic mice. (**A**) Experimental design: Mice on Western diet were subjected to MCAO and treated with vehicle or MSC-EVs (2x10^6^ cell equivalents) immediately after reperfusion. Rosuvastatin (5 mg/kg) was intraperitoneally administered immediately, 24 hours, and 48 hours after MCAO. Neurological deficits were evaluated at 24, 48, and 72 hours using the Clark score, followed by sacrifice at 72 hours post-MCAO (created with https://BioRender.com). (**B**) Neurological deficits using the Clark score assessed at 24, 48, and 72 hours post-MCAO and (**C**) infarct volume and brain edema evaluated by cresyl violet staining at 72 hours post-MCAO. (**D**) Experimental design: Rosuvastatin (5 mg/kg) was intraperitoneally administered once daily for 7 consecutive days to mice on Western diet for 5 weeks. The day after the last rosuvastatin injection, mice were subjected to MCAO, followed by intravenous administration of vehicle or MSC-EVs (2x10^6^ cell equivalents) immediately after reperfusion. Neurological deficits were evaluated at 24, 48, and 72 hours after MCAO, followed by sacrifice at 72 hours (created with https://BioRender.com). (**E**) Neurological deficits at 24, 48, and 72 hours and (**F**) infarct volume and brain edema at 72 hours post-MCAO. Representative cresyl violet stainings are shown (in (**C, F**)). Data were evaluated by one-way ANOVA with LSD post hoc tests (in (**B, C, E, F**)). No significant group differences were noted (n=14-17 mice/group (in (**B, C**)), n=6 mice/group (in (**E, F**))). Scale bars: 1 mm (in (**C, F**)).

### Hyperlipidemia disrupts systemic innate immune balance, which can be restored by DNase-I

Because peripheral monocytes/ macrophages and neutrophils are indispensable for MSC-EV-induced neuroprotection in ischemic stroke models,^7,11^ we hypothesized that hyperlipidemic mice on Western diet may develop a systemic immune profile that renders them unresponsive to MSC-EV treatment. To test this, blood samples of mice fed with normal diet or Western diet for 6 weeks were analyzed by flow cytometry (Figure 3A). Hyperlipidemia significantly reduced the frequencies of macrophages, resting neutrophils, and T cells, while increasing the frequencies of activated and aged neutrophils in the blood (Figure 3B). Although the frequencies of total monocytes and monocyte subsets—including classical Ly6C^high^ inflammatory, Ly6C^int^ intermediate, and non-classical Ly6C^low^ patrolling monocytes—remained unchanged, MHC II expression on monocytes was significantly reduced in hyperlipidemic mice (Figure 3B, Figure S2). Consistent with neutrophil activation leading to the release of neutrophil-derived extracellular DNA traps,^30,31^ circulating cell-free DNA levels were elevated in the blood of hyperlipidemic compared with normolipidemic mice (Figure 3C). These findings indicate that 6 weeks of Western diet feeding predominantly disrupts systemic innate immune homeostasis, characterized by impaired monocyte/macrophage activation and neutrophil overactivation. As degradation of circulating cell-free DNA by DNase-I has been shown to exert anti-inflammatory effects and attenuate immune dysfunction after MCAO,^30^ we next investigated whether DNase-I could rescue systemic innate immune dysregulation in hyperlipidemic mice. Following 6 weeks of Western diet, mice received vehicle (normal saline) or DNase-I, and blood was analyzed 24 hours later by flow cytometry (Figure 3D). Compared with vehicle-treated controls, DNase-I increased frequencies of macrophages, M1-like macrophages, M2-like macrophages, dendritic cells, and activated CD4^+^ T cells, accompanied by a reduction in activated neutrophils (Figure 3E, Figure S3). Although total monocytes and monocyte subset frequencies were again unchanged, DNase-I promoted phenotypic reprogramming of monocytes, reflected by significantly increased CD38 expression and a trend toward enhanced MHC II expression (p=0.0684; Figure 3E, Figure S3). Collectively, these findings demonstrate that DNase-I rebalanced Western diet-induced immune changes by reducing neutrophil activation and restoring monocyte/ macrophage phenotypes.

**Figure 3.**
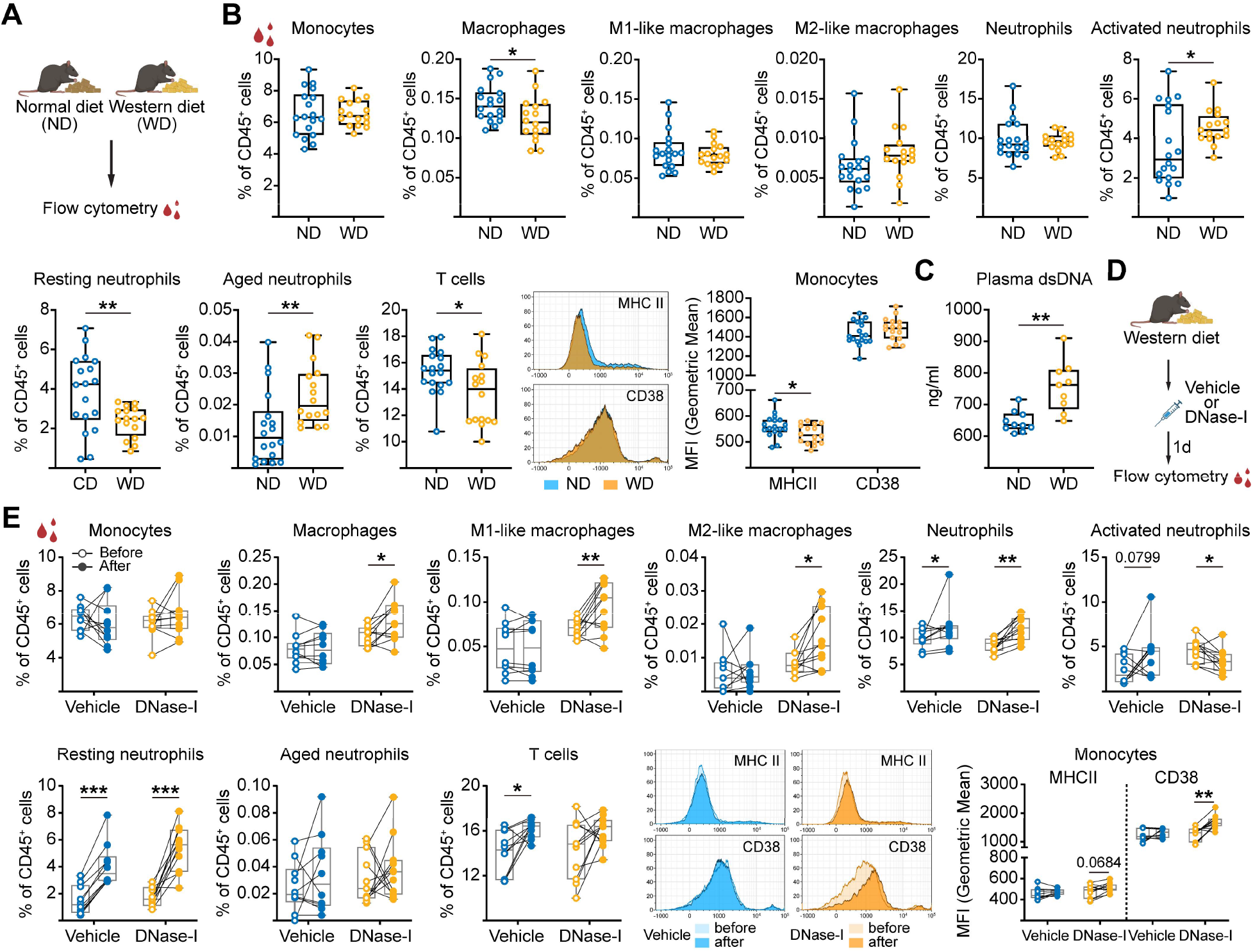
Hyperlipidemia disrupts innate immune balance, which can be restored by DNase-I. (**A**) Experimental design: Blood samples were collected from mice after 6 weeks of normal diet (ND) or Western diet (WD) and analyzed by flow cytometry (created with https://BioRender.com). (**B**) Flow cytometric quantification of leukocyte subsets in blood, including monocytes (CD11b^+^ CD115^+^), macrophages (CD11b^+^ F4/80^+^ Ly6C^-^ MHC II^+^), M1-like macrophages (CD11b^+^ F4/80^+^ Ly6C^-^ MHC II^+^ CD38^+^ CD206^-^), M2-like macrophages (CD11b^+^ F4/80^+^ Ly6C^-^ MHC II^-^ CD38^-^ CD206^+^), neutrophils (Ly6G^+^), activated neutrophils (Ly6G^+^ CXCR2^+^ CD62L^low^), resting neutrophils (Ly6G^+^ CXCR2^+^ CXCR4^-^ CD62L^high^ CD54^low^), aged neutrophils (Ly6G^high^ CXCR4^+^ CD62L^low^), and T cells (CD3e^+^). Representative histograms and corresponding quantification of MHC II and CD38 expression on circulating monocytes. (**C**) Quantification of cell-free double-strand DNA (dsDNA) levels in the blood of mice after 6 weeks of normal diet or Western diet. (**D**) Experimental design: Hyperlipidemic mice on Western diet for 6 weeks received vehicle or DNase-I. Blood samples were collected 24 hours later for flow cytometry (created with https://BioRender.com). (**E**) Quantification of leukocyte subsets in blood, including monocytes, macrophages, M1-like macrophages, M2-like macrophages, neutrophils, activated neutrophils, resting neutrophils, aged neutrophils, and T cells (as above). Representative histograms and corresponding quantification of MHC II and CD38 expression on circulating monocytes. Data were evaluated by unpaired t tests or Mann-Whitney U tests (in (**B, C**)) or paired t tests (in (**E**)). *p<0.05, **p<0.01, ***p<0.001 (n=16-18 mice/group (in (**B**)), n=9-10 mice/group (in (**C**)), n=10-11 mice/group (in (**E**)); each pair represents one individual mouse).

### DNase-I preconditioning salvages ischemic neuroprotection by MSC-EVs

We next investigated whether DNase-I could rescue the neuroprotective efficacy of MSC-EVs in hyperlipidemic MCAO mice. After 6 weeks of Western diet, mice received vehicle or DNase-I, followed by MCAO 24 hours later and MSC-EV treatment immediately after reperfusion onset (Figure 4A). In DNase-I-preconditioned mice, MSC-EV treatment significantly reduced neurological deficits, infarct volume and brain edema (Figure 4B, C). MSC-EVs also showed a trend toward reducing the density of DNA-fragmented TUNEL^+^/ NeuN^+^ injured neurons in the ischemic striatum (p=0.0743; Figure D). In addition, MSC-EVs significantly reduced the density of brain-infiltrated neutrophils (Figure 4E), but did not significantly influence total leukocyte infiltration (Figure S4A), ICAM-1 expression, a leukocyte adhesion molecule on cerebral microvessels (Figure S4B), GPIbα^+^ microvascular thrombosis (Figure S4C), or IgG extravasation as a marker of blood-brain barrier permeability (Figure S4D). DNase-I pretreatment alone or MSC-EV treatment alone had no significant effects on any of the readouts examined (Figure 4B-E, Figure S4). Flow cytometric analysis furthermore revealed that MSC-EVs significantly reduced brain-infiltrated neutrophils, activated neutrophils, resting neutrophils, N1-like neutrophils, and N2-like neutrophils in DNase-I-preconditioned hyperlipidemic mice (Figure 5). In addition, trends toward reduced overall leukocytes, monocytes, intermediate monocytes, inflammatory monocytes, M1-like macrophages, and activated CD4^+^ T cells were noted (Figure 5). Neither DNase-I pretreatment alone nor MSC-EV treatment alone significantly influenced post-ischemic brain leukocyte infiltrates (Figure 5). Likewise, no significant changes in peripheral blood leukocyte populations were detected in response to either treatment alone or their combination (Figure S5). Together, these findings indicate that DNase-I preconditioning restores MSC-EV-induced neuroprotection and immunomodulatory activity in hyperlipidemic mice post-MCAO.

**Figure 4.**
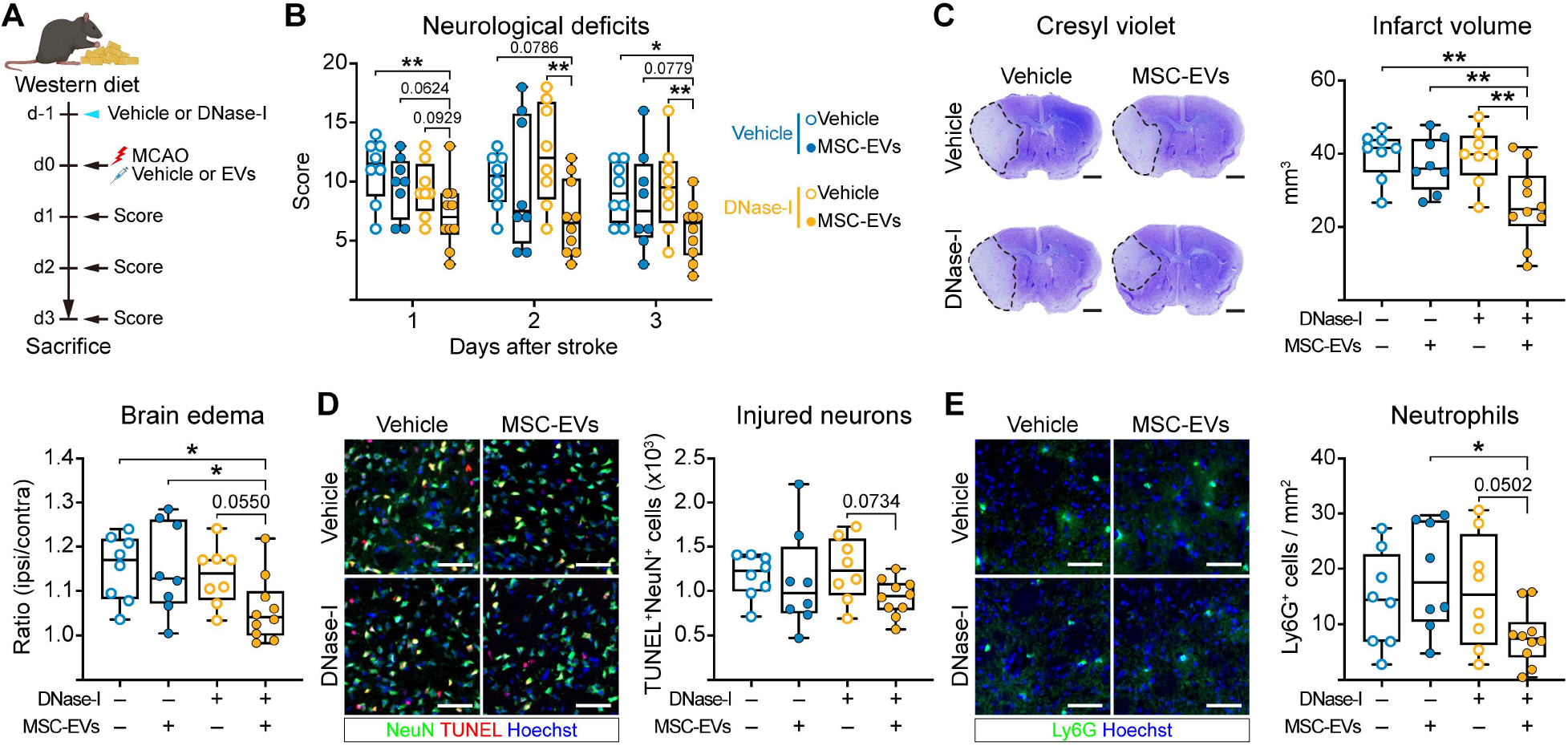
DNase-I preconditioning salvages MSC-EV-induced ischemic neuroprotection in hyperlipidemic mice. (**A**) Experimental design: Vehicle or DNase-I was administered to hyperlipidemic mice on Western diet 24 hours before MCAO. Vehicle or MSC-EVs (2x10^6^ cell equivalents) were intravenously administered immediately after reperfusion. Neurological deficits were evaluated at 24, 48, and 72 hours post-MCAO, followed by animal sacrifice at 72 hours for histochemical analyses (created with https://BioRender.com). (**B**) Neurological deficits assessed using the Clark score at 24, 48, and 72 hours post-MCAO. (**C**) Infarct volume and brain edema evaluated by cresyl violet staining, (**D**) neuronal injury in the ischemic striatum assessed by TUNEL/ NeuN immunolabeling, and (**E**) density of brain-infiltrated neutrophils in the ischemic striatum at 72 hours post-MCAO. Representative cresyl violet stainings (in (**C**)) and immunohistochemistry images (in (**D**, **E**)) are shown. Data were compared by two-way ANOVA with LSD post hoc tests (in (**B-E**)). *p<0.05, **p<0.01 (n=8-10 mice/group). Scale bars: 1 mm (in (**C**)), 50 μm (in (**D**, **E**)).

**Figure 5.**
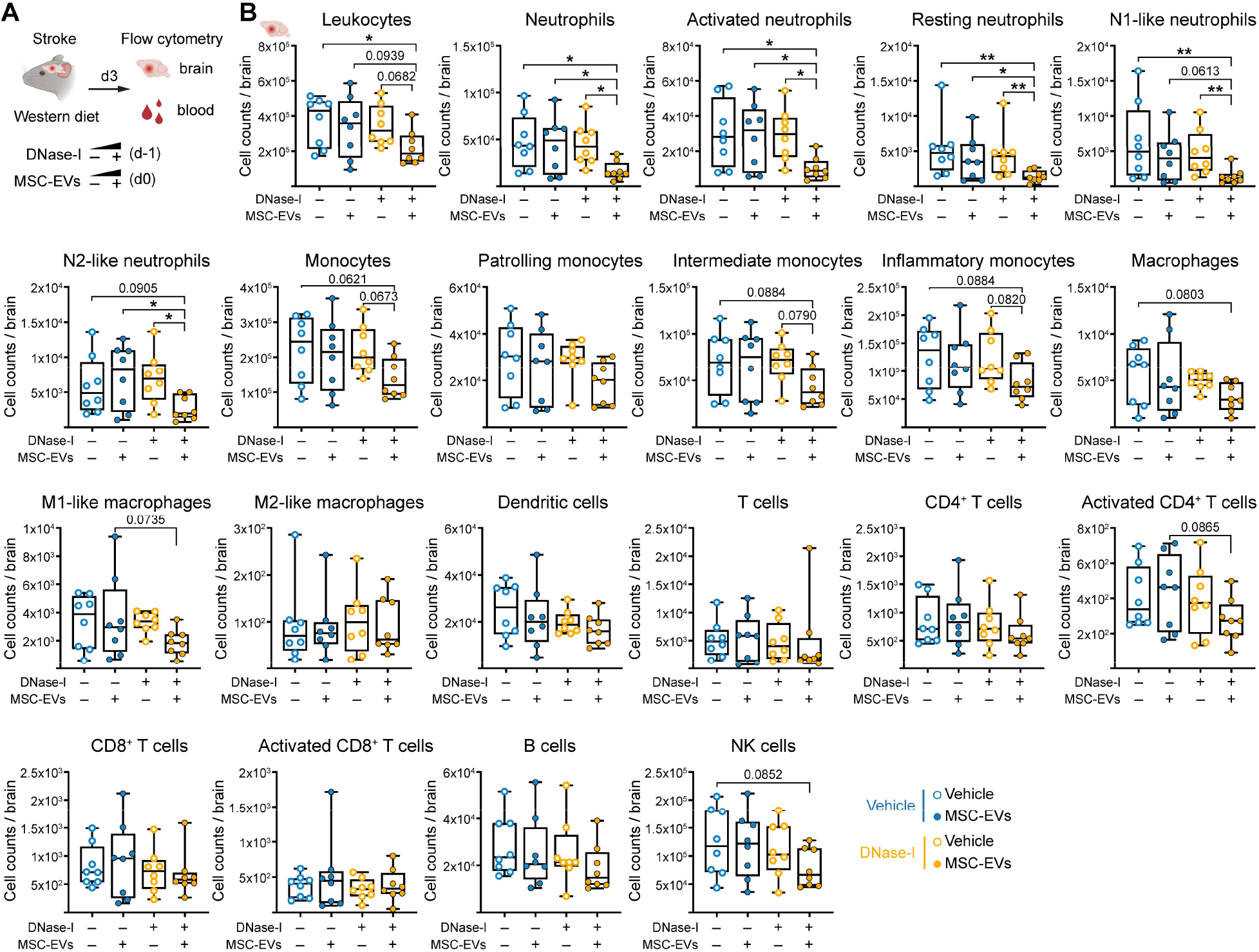
DNase-I preconditioning restores MSC-EV-induced prevention of brain leukocyte entry in hyperlipidemic mice. (**A**) Experimental design: Vehicle or DNase-I was administered to hyperlipidemic mice on Western diet 24 hours before MCAO, followed by intravenous administration of vehicle or MSC-EVs (2x10^6^ cell equivalents) immediately after reperfusion. Ischemic brain tissue and peripheral blood were collected 72 hours after MCAO for flow cytometry (created with https://BioRender.com). (**B**) Quantification of leukocyte subsets in the ischemic brain, including total leukocytes, neutrophils, activated neutrophils, resting neutrophils, N1-like neutrophils (Ly6G^+^ CD170^low^ CD54^+^ PD-L1^-^), N2-like neutrophils (Ly6G^+^ CD170^high^ PD-L1^+^), monocytes, patrolling monocytes (CD11b^+^ CD115^+^ Ly6C^low^), intermediate monocytes (CD11b^+^ CD115^+^ Ly6C^int^), inflammatory monocytes (CD11b^+^ CD115^+^ Ly6C^high^), macrophages, M1-like macrophages, M2-like macrophages, dendritic cells (F4/80^-^ CD11c^+^ MHC II^+^), T cells, CD4^+^ T cells (CD3e^+^ CD4^+^), activated CD4^+^ T cells (CD3e^+^ CD4^+^ CD69^+^), CD8^+^ T cells (CD3e^+^ CD8^+^), activated CD8^+^ T cells (CD3e^+^ CD8^+^ CD69^+^), B cells (CD3e^-^B220^+^), and NK cells (CD3e^-^ B220^-^ NK-1.1^+^). Data were compared by two-way ANOVA with LSD post hoc tests (in (**B**)). *p<0.05, **p<0.01 (n=8 mice/group).

### Post-ischemic DNase-I treatment does not reestablish MSC-EV-induced neuroprotection

Circulating cell-free DNA can associate with the surface of EVs, thereby increasing their net negative charge, promoting aggregation, and ultimately influencing their biological activity in recipient cells.^32^ To explore whether prevention of cell-free DNA accumulation on EVs restores the neuroprotective effects of MSC-EVs, we next performed experiments, in which we administered DNase-I immediately after MCAO shortly before MSC-EV delivery (Figure 6A). By this approach, cell-free DNA would be degraded at the time of MSC-EV administration, whereas Western diet-induced systemic immune alterations would remain largely unchanged in the early stroke phase. Unlike DNase-I preconditioning, DNase-I treatment immediately after reperfusion induced ischemic neuroprotection, reducing neurological deficits, infarct volume, brain edema, brain leukocyte infiltrates—specifically neutrophil infiltrates—and microvascular thrombosis, whereas neuronal injury, ICAM-1 abundance, or blood-brain barrier permeability remained unchanged (Figure 6B-E, Figure S6). Importantly, post-ischemic DNase-I treatment did not reestablish the neuroprotective effects of MSC-EVs in hyperlipidemic mice in any readout examined (Figure 6B-E, Figure S6).

**Figure 6.**
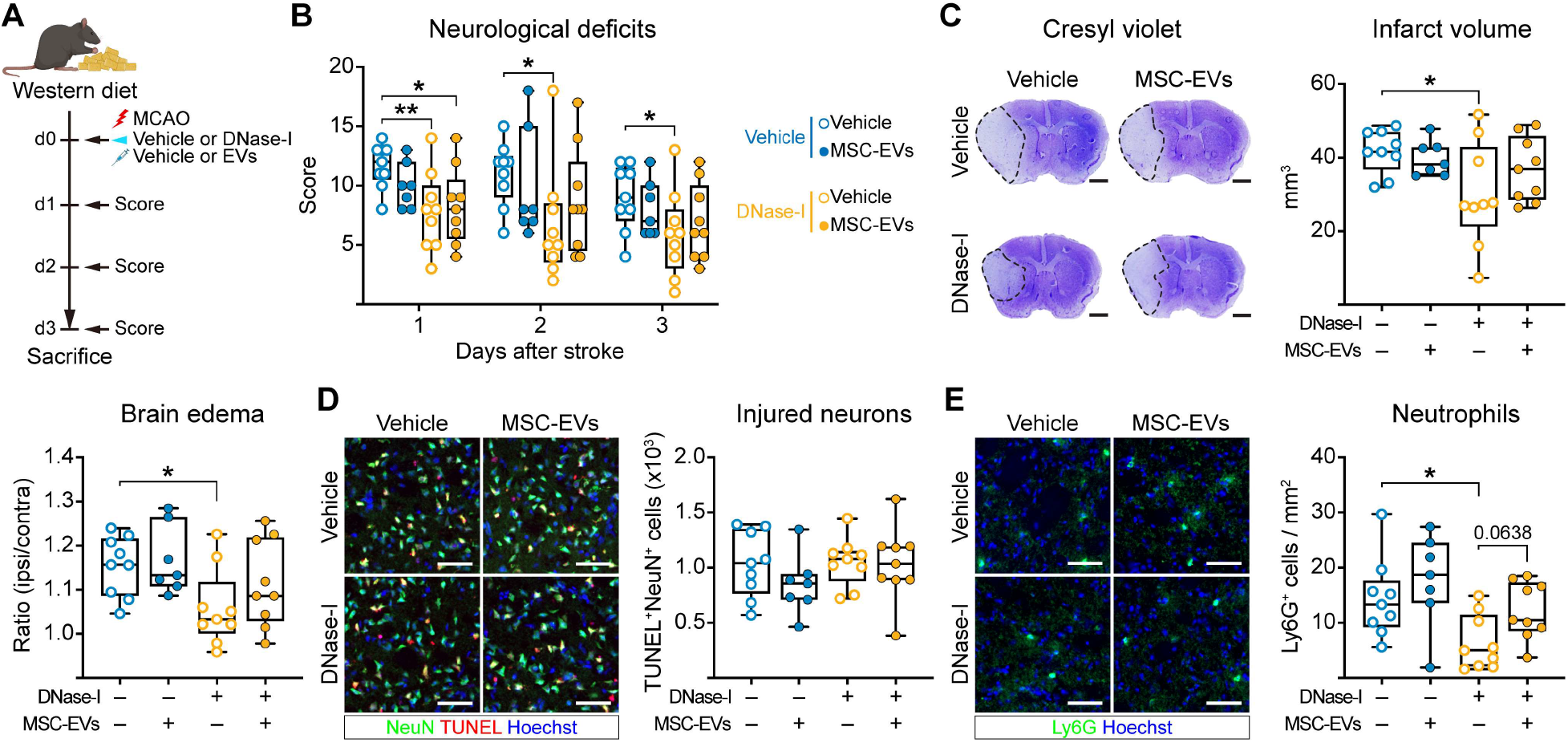
DNase-I administration after MCAO fails to reestablish MSC-EV-induced ischemic neuroprotection in hyperlipidemic mice. (**A**) Experimental design: Vehicle or DNase-I was administered to hyperlipidemic mice on Western diet immediately after MCAO, followed by vehicle or MSC-EV (2x10^6^ cell equivalents) treatment. Neurological deficits were evaluated at 24, 48, and 72 hours post-MCAO. Animals were sacrificed at 72 hours post-MCAO for histochemical analyses (created with https://BioRender.com). (**B**) Neurological deficits assessed by Clark score at 24, 48, and 72 hours after MCAO. (**C**) Infarct volume and brain edema evaluated by cresyl violet staining, (**D**) neuronal injury in the ischemic striatum assessed by TUNEL/ NeuN immunolabeling, and (**E**) density of brain-infiltrated neutrophils in the ischemic striatum at 72 hours post-MCAO. Representative cresyl violet stainings (in (**C**)) and immunohistochemistry images (in (**D**, **E**)) are shown. Data were compared by two-way ANOVA with LSD post hoc tests (in (**B-E**)). *p<0.05 (n=7-9 mice/group). Scale bars: 1 mm (in (**C**)), 50 μm (in (**D**, **E**)).

### Monocytes/ macrophages and neutrophils are indispensable for MSC-EV neuroprotection in hyperlipidemic mice

To further explore the role of monocytes/ macrophages and neutrophils in restoring MSC-EV responsiveness, we preconditioned hyperlipidemic mice with DNase-I and depleted monocytes/ macrophages or neutrophils immediately thereafter using clodronate liposomes or anti-Ly6G antibody, respectively (Figure 7A). In hyperlipidemic mice exposed to combined DNase-I preconditioning and monocyte/ macrophage depletion, MSC-EV treatment not only failed to induce neuroprotection compared with vehicle treatment, but significantly increased neurological deficits, infarct volume, brain edema, and brain leukocyte infiltrates, specifically neutrophil infiltrates, as well as ICAM-1 expression on cerebral microvessels in the ischemic striatum (Figure 7B-E, Figure S7). In mice receiving combined DNase-I preconditioning and neutrophil depletion, on the other hand, MSC-EV treatment failed to influence stroke outcomes (Figure 7B-E, Figure S7). The consequences of monocyte/ macrophage or neutrophil depletion are consistent with previous findings of our group in normolipidemic MCAO mice, in which neutrophil depletion abolished MSC-EV-induced neuroprotection whereas monocyte/ macrophage depletion turned neuroprotection by MSC-EVs into the exacerbation of ischemic brain injury.^7,11^ Collectively, these findings identify monocytes/ macrophages and neutrophils as critical mediators of MSC-EV-induced neuroprotection in ischemic stroke.

**Figure 7.**
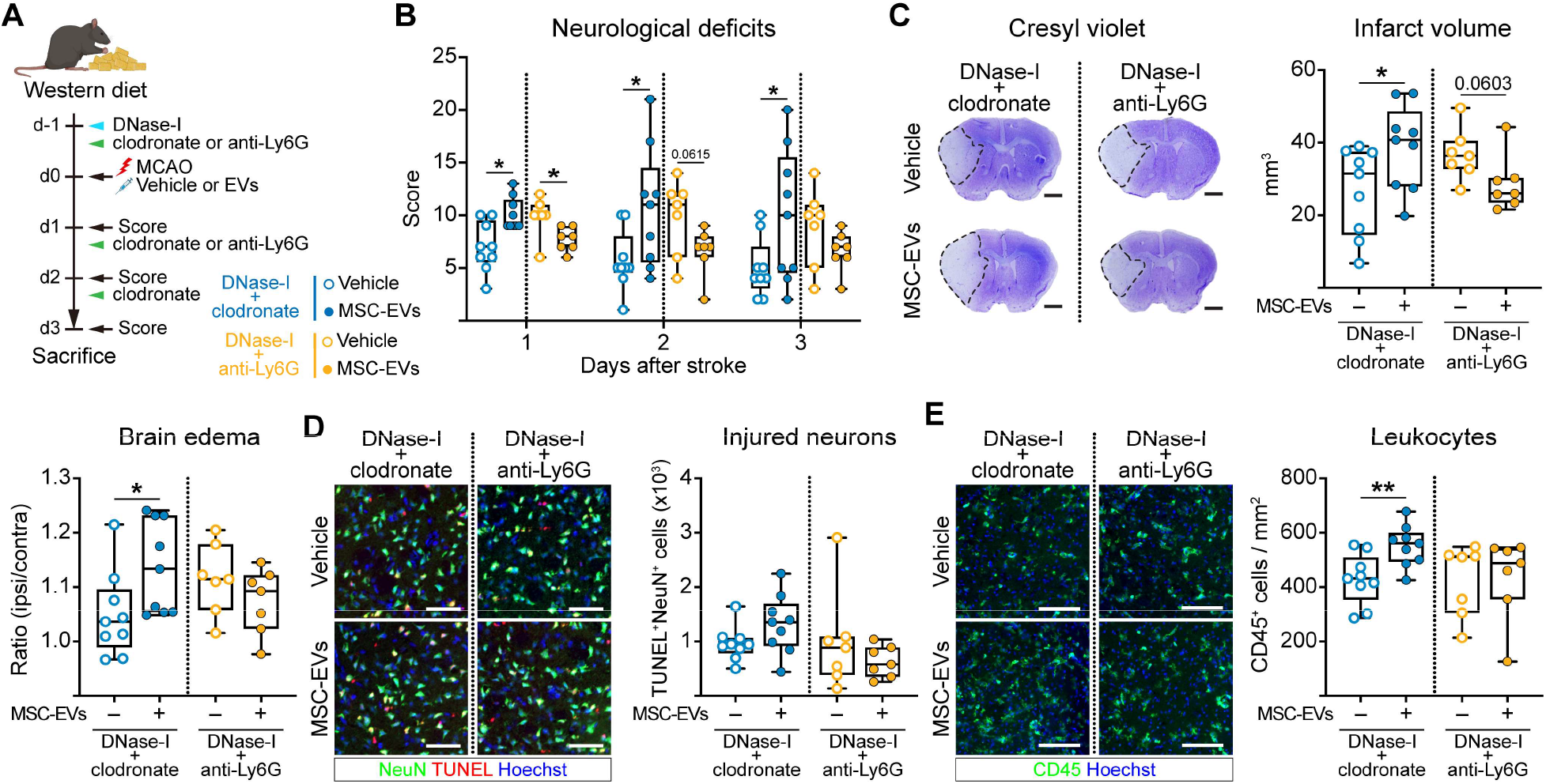
Monocyte/ macrophage or neutrophil depletion abolishes MSC-EV-induced neuroprotection in DNase-I-preconditioned hyperlipidemic mice. (**A**) Experimental design: Vehicle or DNase-I was administered to hyperlipidemic mice on Western diet 24 hours before MCAO, followed by the depletion of monocytes/ macrophages or neutrophils using clodronate liposomes or anti-Ly6G antibody, respectively. Vehicle or MSC-EVs (2x10^6^ cell equivalents) were intravenously administered immediately after reperfusion onset (created with https://BioRender.com). (**B**) Neurological deficits assessed by Clark score at 24, 48, and 72 hours post-MCAO. (**C**) Infarct volume and brain edema evaluated by cresyl violet staining, (**D**) neuronal injury in the ischemic striatum assessed by TUNEL/ NeuN immunolabeling, and (**E**) density of brain-infiltrated leukocytes in the ischemic striatum at 72 hours post-MCAO. Representative cresyl violet stainings (in (**C**)) and immunohistochemistry images (in (**D, E**)) are shown. Data were compared by unpaired t tests or Mann-Whitney U tests (in (**B-E**)). *p<0.05, **p<0.01 (n=7-9 mice/group). Scale bars: 1 mm (in (**C**)), 50 μm (in (**D**), 100 μm (in (**E**)).

## Discussion

We herein demonstrate that MSC-EVs—despite their confirmed neuroprotective efficacy in young and aged normolipidemic mice and rats^6-9^—fail to induce neuroprotection after MCAO in hyperlipidemic mice on Western diet. Loss of MSC-EV efficacy was associated with profound innate immune dysregulation characterized by reduced monocyte/ macrophage activation, increased neutrophil activation and elevated circulating cell-free DNA levels in the blood. Dose escalation of MSC-EVs or add-on treatment with the lipid-lowering HMG-CoA reductase inhibitor rosuvastatin did not restore MSC-EV-induced neuroprotection. DNase-I administration before MCAO (as a stimulus to rebalance immune dysfunction), but not DNase-I administration after MCAO (as a tool to degrade cell-free DNA released after the stroke) restored MSC-EV-induced neuroprotection and immunomodulation. The recuperation of neuroprotection was closely linked to the restitution of monocyte/ macrophage and neutrophil function, as shown in monocyte/ macrophage and neutrophil depletion studies. These findings identify host innate immune function as a key determinant of MSC-EV efficacy in hyperlipidemic mice.

The failure of neuroprotectants in clinical translation has emphasized the importance of replicating clinically relevant risk factors and diseases in experimental stroke models.^33,34^ Although MSCs and MSC-EVs consistently induced ischemic neuroprotection in otherwise healthy young and aged mice through monocyte/ macrophage- and neutrophil-dependent modes of action,^6-8,11^ the therapeutic efficacy of MSC-EVs may profoundly be altered in vascular risk contexts, as the present study shows. When administered over six weeks, the cholesterol-rich Western diet applied in this study induces fatty streaks in cerebral microvessels, which predispose the brain post-MCAO to exacerbated blood-brain barrier breakdown^13^ and neuroinflammation.^14^ Serum cholesterol levels are increased ∼4-fold in hyperlipidemic C57BL/6j mice on Western diet.^13^ In addition, plasma glucose levels are increased, as the present study shows. It has previously been demonstrated in rats with streptozotocin-induced type-I diabetes that treatment with bone marrow-derived MSCs did not improve neurological recovery after MCAO, but increased mortality, blood-brain barrier leakage, and brain hemorrhage formation.^35-37^ In histochemical studies, neointima formation and arteriole narrowing were exacerbated by bone marrow-derived MSCs in type-1 diabetic rats.^35,36^ These abnormalities were attributed to increased angiogenin expression in the brain and brain-supplying arteries of these rats.^35^ In contrast to type-1 diabetes, MSCs enhanced neurological recovery, reduced blood–brain barrier leakage and reduced brain hemorrhage formation after MCAO in type-2 diabetic rats.^38^ MSCs and MSC-EVs also increased neurological recovery, blood-brain barrier integrity, white matter remodeling and axonal outgrowth after MCAO in type-2 diabetic rats, even when obtained from type-2 diabetic rats.^39,40^ The underlying reasons for the compromised MSC and MSC-EV responses in type-1, but not type-2 diabetes remained unclear. Unlike in type-1 diabetic rats, MSC-EVs did not deteriorate stroke outcome in hyperlipidemic mice on Western diet, but abolished the beneficial effects on stroke outcome.

Hyperlipidemia promotes neutrophil activation accompanied by elevated levels of circulating neutrophil extracellular DNA traps (NETs).^16,17^ NETs causatively contribute to poststroke immunosuppression^30^ and microvascular thromboinflammation.^41^ Of note, the loss of MSC-EV actions in hyperlipidemic mice cannot be explained solely by direct EV interactions with circulating lipoproteins or extracellular DNA. In fact, MSC-EV dose-escalation failed to restore neuroprotection, as did lipid-lowering with rosuvastatin or cell-free DNA degradation by DNase-I administration after MCAO, which removed DNA released on occasion of the stroke. Instead, the restitution of MSC-EV-induced neuroprotection required prior immune reprogramming, which we achieved by pre-ischemic preconditioning with DNase-I.^30,31^ DNase-I preconditioning rebalanced the Western diet-induced immune changes, restoring monocyte/ macrophage and neutrophil numbers and activation, and reestablished an immune microenvironment permissive to MSC-EV-induced immunomodulation, as our flow cytometry studies showed. As a result, MSC-EVs successfully reduced infiltrated neutrophils, activated neutrophils, N1-like neutrophils, and N2-like neutrophils in the brains of DNase-I-preconditioned hyperlipidemic MCAO mice. Notably, the depletion of either monocytes/ macrophages or neutrophils abolished the neuroprotective effects of MSC-EVs, confirming earlier findings of our group in normolipidemic mice pinpointing the importance of both myeloid cell entities in MSC-EV neuroprotection.^7,10,11^

Although DNase-I administration after MCAO did not restore MSC-EV responsiveness, post-ischemic DNase-I administration alone induced neuroprotection in hyperlipidemic MCAO mice, reducing neurological deficits, infarct volume, brain edema, brain neutrophil infiltrates, and microvascular thrombosis. This observation is consistent with previous studies demonstrating that DNase-I improves neurological recovery and decreases infarct volume and microvascular thrombosis post-MCAO in otherwise healthy mice and rats.^30,41,42^ Although DNase-I itself represents a promising neuroprotective strategy, two limitations should be considered. First, cell-free DNA is rapidly released by neutrophils after ischemic stroke, leaving a short time-window for DNase-I administration.^30,31^ Secondly, DNase-I administration has been reported to increase hemorrhagic risk after delayed (72 hours post-MCAO) but not early (immediately after MCAO) post-ischemic administration.^27,42^ Both aspects imply that DNase-I treatment timing is critical for the efficacy and safety of this drug. That post-ischemic DNase-I administration induces neuroprotection in hyperlipidemic MCAO mice, has to the best of our knowledge not been shown. Future studies should determine immune-priming approaches other than DNase-I, with the idea of reestablishing permissive environments that can successfully be targeted under vascular risk profile conditions.

Our findings have implications for the clinical translation of MSC-EVs. With their potent effects in ischemic stroke models, MSC-EVs are rapidly advancing towards clinical trials. The immunomodulatory mode-of-action of MSC-EVs opens ways for measuring immune responses to MSC-EVs in stroke patients in early randomized controlled phase IIa trials. The assessment of immune responses in the blood will open promising perspectives to prove the concept of MSC-EV-induced immunomodulation in clinical settings. Such studies should carefully check if immune responses are altered in stroke patients with vascular risk profiles. Evaluating patients at risk will increase the chances of successful clinical translation of MSC-EV therapies. Biomarker-based treatment monitoring will lift MSC-EV therapies to the next level.

## Acknowledgments

We thank the Imaging Center Essen (IMCES) at the Faculty of Medicine, University of Duisburg-Essen, Germany, for providing imaging support.

## Sources of Funding

This work was supported by German Research Foundation (DFG) grants 514990328, 449437943 (C6, within TRR332 “Neutrophils: Origin, fate, and function”), and 405358801/428817542 (A4, within FOR2879 “ImmunoStroke”), by the Romanian Executive Agency for Higher Education, Research, Development and Innovation Funding (UEFISCDI) grant PN-IV-P1-PCE-2023-1289 and by the European Union’s “National Recovery and Resilience Plan” through the project “Targeting macrophages/monocytes in the aged ischemic brain by pharmacological, genetic, and cell-based tools” (project no. 760058) (all to DMH). DMH also received funding via E.U. ERA-NET NEURON grant 101168752 (SECRET) and German Federal Ministry of Research, Technology, and Space (BMFTR) – NIH - Collaborative Research grant 01GQ2405B (TopoVess).

## Supplemental Material

Supplemental Methods

Figures S1-S7

Tables S1, S2

References 43, 44

## Notes

### Competing Interest Statement

The authors have declared no competing interest.

## References

1. Albers GW, Marks MP, Kemp S, Christensen S, Tsai JP, Ortega-Gutierrez S, McTaggart RA, Torbey MT, Kim-Tenser M, Leslie-Mazwi T, et al. Thrombectomy for Stroke at 6 to 16 Hours with Selection by Perfusion Imaging. N Engl J Med. 2018;378:708–718. doi: doi:10.1056/NEJMoa1713973

2. Campbell BCV, Mitchell PJ, Churilov L, Yassi N, Kleinig TJ, Dowling RJ, Yan B, Bush SJ, Dewey HM, Thijs V, et al. Tenecteplase versus Alteplase before Thrombectomy for Ischemic Stroke. N Engl J Med. 2018;378:1573–1582. doi: doi:10.1056/NEJMoa1716405

3. Neumann J, Riek-Burchardt M, Herz J, Doeppner TR, Konig R, Hutten H, Etemire E, Mann L, Klingberg A, Fischer T, et al. Very-late-antigen-4 (VLA-4)-mediated brain invasion by neutrophils leads to interactions with microglia, increased ischemic injury and impaired behavior in experimental stroke. Acta Neuropathol. 2015;129:259–277. doi: 10.1007/s00401-014-1355-2

4. Perez-de-Puig I, Miró-Mur F, Ferrer-Ferrer M, Gelpi E, Pedragosa J, Justicia C, Urra X, Chamorro A, Planas AM. Neutrophil recruitment to the brain in mouse and human ischemic stroke. Acta Neuropathol. 2015;129:239–257. doi: 10.1007/s00401-014-1381-0

5. Hermann DM, Peruzzotti-Jametti L, Giebel B, Pluchino S. Extracellular vesicles set the stage for brain plasticity and recovery by multimodal signalling. Brain. 2024;147:372–389. doi: 10.1093/brain/awad332

6. Doeppner TR, Herz J, Gorgens A, Schlechter J, Ludwig AK, Radtke S, de Miroschedji K, Horn PA, Giebel B, Hermann DM. Extracellular Vesicles Improve Post-Stroke Neuroregeneration and Prevent Postischemic Immunosuppression. Stem Cells Transl Med. 2015;4:1131–1143. doi: 10.5966/sctm.2015-0078

7. Wang C, Börger V, Sardari M, Murke F, Skuljec J, Pul R, Hagemann N, Dzyubenko E, Dittrich R, Gregorius J, et al. Mesenchymal Stromal Cell-Derived Small Extracellular Vesicles Induce Ischemic Neuroprotection by Modulating Leukocytes and Specifically Neutrophils. Stroke. 2020;51:1825–1834. doi: 10.1161/strokeaha.119.028012

8. Wang C, Börger V, Yusuf AM, Tertel T, Stambouli O, Murke F, Freund N, Kleinschnitz C, Herz J, Gunzer M, et al. Postischemic Neuroprotection Associated With Anti-Inflammatory Effects by Mesenchymal Stromal Cell-Derived Small Extracellular Vesicles in Aged Mice. Stroke. 2022;53:e14–e18. doi: doi:10.1161/STROKEAHA.121.035821

9. Xin H, Li Y, Cui Y, Yang JJ, Zhang ZG, Chopp M. Systemic administration of exosomes released from mesenchymal stromal cells promote functional recovery and neurovascular plasticity after stroke in rats. J Cereb Blood Flow Metab. 2013;33:1711–1715. doi: 10.1038/jcbfm.2013.152

10. Gregorius J, Wang C, Stambouli O, Hussner T, Qi Y, Tertel T, Börger V, Yusuf AM, Hagemann N, Yin D, et al. Small extracellular vesicles obtained from hypoxic mesenchymal stromal cells have unique characteristics that promote cerebral angiogenesis, brain remodeling and neurological recovery after focal cerebral ischemia in mice. Basic Res Cardiol. 2021;116:40. doi: 10.1007/s00395-021-00881-9

11. Wang C, Zhang Y, Tertel T, Mouloud Y, Liu X, Hagemann N, Mohamud Yusuf A, Gronewold J, Strecker J-K, Popa-Wagner A, et al. Monocytes shape the neuroprotective and immunomodulatory effects of mesenchymal stromal cell-derived extracellular vesicles. bioRxiv. 2026:2026.2006.2001.727369. doi: 10.64898/2026.06.01.727369

12. Sacco RL, Diener H-C, Yusuf S, Cotton D, Ôunpuu S, Lawton WA, Palesch Y, Martin RH, Albers GW, Bath P, et al. Aspirin and Extended-Release Dipyridamole versus Clopidogrel for Recurrent Stroke. N Engl J Med. 2008;359:1238–1251. doi: doi:10.1056/NEJMoa0805002

13. ElAli A, Doeppner TR, Zechariah A, Hermann DM. Increased blood-brain barrier permeability and brain edema after focal cerebral ischemia induced by hyperlipidemia: role of lipid peroxidation and calpain-1/2, matrix metalloproteinase-2/9, and RhoA overactivation. Stroke. 2011;42:3238–3244. doi: 10.1161/strokeaha.111.615559

14. Herz J, Hagen SI, Bergmüller E, Sabellek P, Göthert JR, Buer J, Hansen W, Hermann DM, Doeppner TR. Exacerbation of ischemic brain injury in hypercholesterolemic mice is associated with pronounced changes in peripheral and cerebral immune responses. Neurobiol Dis. 2014;62:456–468. doi: 10.1016/j.nbd.2013.10.022

15. Christ A, Günther P, Lauterbach MAR, Duewell P, Biswas D, Pelka K, Scholz CJ, Oosting M, Haendler K, Baßler K, et al. Western Diet Triggers NLRP3-Dependent Innate Immune Reprogramming. Cell. 2018;172:162–175.e114. doi: 10.1016/j.cell.2017.12.013

16. Dhawan UK, Bhattacharya P, Narayanan S, Manickam V, Aggarwal A, Subramanian M. Hypercholesterolemia Impairs Clearance of Neutrophil Extracellular Traps and Promotes Inflammation and Atherosclerotic Plaque Progression. Arterioscler Thromb Vasc Biol. 2021;41:2598–2615. doi: doi:10.1161/ATVBAHA.120.316389

17. Drechsler M, Megens RTA, Zandvoort Mv, Weber C, Soehnlein O. Hyperlipidemia-Triggered Neutrophilia Promotes Early Atherosclerosis. Circulation. 2010;122:1837–1845. doi: doi:10.1161/CIRCULATIONAHA.110.961714

18. Zechariah A, ElAli A, Hagemann N, Jin F, Doeppner TR, Helfrich I, Mies G, Hermann DM. Hyperlipidemia Attenuates Vascular Endothelial Growth Factor–Induced Angiogenesis, Impairs Cerebral Blood Flow, and Disturbs Stroke Recovery via Decreased Pericyte Coverage of Brain Endothelial Cells. Arterioscler Thromb Vasc Biol. 2013;33:1561–1567. doi: doi:10.1161/ATVBAHA.112.300749

19. Tuz AA, Hoerenbaum N, Ulusoy Ö, Ahmadi A, Gerlach A, Beer A, Kraus A, Hasenberg A, Hagemann N, Hermann DM, et al. Hypercholesterolemia triggers innate immune imbalance and transforms brain infarcts after ischemic stroke. Front Immunol. 2025;15. doi: 10.3389/fimmu.2024.1502346

20. Herz J, Sabellek P, Lane TE, Gunzer M, Hermann DM, Doeppner TR. Role of Neutrophils in Exacerbation of Brain Injury After Focal Cerebral Ischemia in Hyperlipidemic Mice. Stroke. 2015;46:2916–2925. doi: 10.1161/strokeaha.115.010620

21. Yin D, Wang C, Qi Y, Wang Y-C, Hagemann N, Mohamud Yusuf A, Dzyubenko E, Kaltwasser B, Tertel T, Giebel B, et al. Neural precursor cell delivery induces acute post-ischemic cerebroprotection, but fails to promote long-term stroke recovery in hyperlipidemic mice due to mechanisms that include pro-inflammatory responses associated with brain hemorrhages. J Neuroinflamm. 2023;20:210. doi: 10.1186/s12974-023-02894-8

22. Percie du Sert N, Hurst V, Ahluwalia A, Alam S, Avey MT, Baker M, Browne WJ, Clark A, Cuthill IC, Dirnagl U, et al. The ARRIVE guidelines 2.0: Updated guidelines for reporting animal research. J Cereb Blood Flow Metab. 2020;40:1769–1777. doi: 10.1371/journal.pbio.3000410

23. Labusek N, Mouloud Y, Köster C, Diesterbeck E, Tertel T, Wiek C, Hanenberg H, Horn PA, Felderhoff-Müser U, Bendix I, et al. Extracellular vesicles from immortalized mesenchymal stromal cells protect against neonatal hypoxic-ischemic brain injury. Inflamm Regen. 2023;43:24. doi: 10.1186/s41232-023-00274-6

24. Labusek N, Ghari P, Mouloud Y, Köster C, Diesterbeck E, Hadamitzky M, Felderhoff-Müser U, Bendix I, Giebel B, Herz J. Hypothermia combined with extracellular vesicles from clonally expanded immortalized mesenchymal stromal cells improves neurodevelopmental impairment in neonatal hypoxic-ischemic brain injury. J Neuroinflamm. 2023;20:280. doi: 10.1186/s12974-023-02961-0

25. Théry C, Witwer KW, Aikawa E, Alcaraz MJ, Anderson JD, Andriantsitohaina R, Antoniou A, Arab T, Archer F, Atkin-Smith GK, et al. Minimal information for studies of extracellular vesicles 2018 (MISEV2018): a position statement of the International Society for Extracellular Vesicles and update of the MISEV2014 guidelines. J Extracell Vesicles. 2018;7:1535750. doi: 10.1080/20013078.2018.1535750

26. Kilic Ü, Bassetti CL, Kilic E, Xing H, Wang Z, Hermann DM. Post-ischemic delivery of the 3-hydroxy-3-methylglutaryl coenzyme A reductase inhibitor rosuvastatin protects against focal cerebral ischemia in mice via inhibition of extracellular-regulated kinase-1/-2. Neuroscience. 2005;134:901–906. doi: 10.1016/j.neuroscience.2005.04.063

27. Yin D, Wang C, Singh V, Tuz AA, Doeppner TR, Gunzer M, Hermann DM. Delayed DNase-I Administration but Not Gasdermin-D Inhibition Induces Hemorrhagic Transformation After Transient Focal Cerebral Ischemia in Mice. Stroke. 2024;55:e297–e299. doi: doi:10.1161/STROKEAHA.124.047862

28. Busatto S, Yang Y, Iannotta D, Davidovich I, Talmon Y, Wolfram J. Considerations for extracellular vesicle and lipoprotein interactions in cell culture assays. J Extracell Vesicles. 2022;11:e12202. doi: 10.1002/jev2.12202

29. Ghebosu RE, Pendiuk Goncalves J, Wolfram J. Extracellular Vesicle and Lipoprotein Interactions. Nano Letters. 2024;24:1-8. doi: 10.1021/acs.nanolett.3c03579

30. Tuz AA, Ghosh S, Karsch L, Ttoouli D, Sata SP, Ulusoy Ö, Kraus A, Hoerenbaum N, Wolf J-N, Lohmann S, et al. Stroke and myocardial infarction induce neutrophil extracellular trap release disrupting lymphoid organ structure and immunoglobulin secretion. Nature Cardiovascular Research. 2024;3:525–540. doi: 10.1038/s44161-024-00462-8

31. Roth S, Wernsdorf SR, Liesz A. The role of circulating cell-free DNA as an inflammatory mediator after stroke. Semin Immunopathol. 2023;45:411–425. doi: 10.1007/s00281-023-00993-5

32. Shelke GV, Jang SC, Yin Y, Lässer C, Lötvall J. Human mast cells release extracellular vesicle-associated DNA. Matters. 2016;2:e201602000034.

33. Shi L, Rocha M, Leak RK, Zhao J, Bhatia TN, Mu H, Wei Z, Yu F, Weiner SL, Ma F, et al. A new era for stroke therapy: Integrating neurovascular protection with optimal reperfusion. J Cereb Blood Flow Metab. 2018;38:2073–2091. doi: 10.1177/0271678x18798162

34. Boltze J, Fisher M. Cytoprotection Concepts for Ischemic Stroke in the Recanalization Era. Advanced Science. 2026;13:e17043. doi: 10.1002/advs.202517043

35. Chen J, Ye X, Yan T, Zhang C, Yang X-P, Cui X, Cui Y, Zacharek A, Roberts C, Liu X, et al. Adverse Effects of Bone Marrow Stromal Cell Treatment of Stroke in Diabetic Rats. Stroke. 2011;42:3551–3558. doi: doi:10.1161/STROKEAHA.111.627174

36. Yan T, Ye X, Chopp M, Zacharek A, Ning R, Venkat P, Roberts C, Lu M, Chen J. Niaspan Attenuates the Adverse Effects of Bone Marrow Stromal Cell Treatment of Stroke in Type One Diabetic Rats. PLoS One. 2013;8:e81199. doi: 10.1371/journal.pone.0081199

37. Mangin G, Cogo A, Moisan A, Bonnin P, Maïer B, Kubis N, obotRC. Intravenous Administration of Human Adipose Derived-Mesenchymal Stem Cells Is Not Efficient in Diabetic or Hypertensive Mice Subjected to Focal Cerebral Ischemia. Front Neurosci. 2019;Volume 13 - 2019. doi: 10.3389/fnins.2019.00718

38. Ding G, Chen J, Chopp M, Li L, Yan T, Li Q, Cui C, Davarani SPN, Jiang Q. Cell Treatment for Stroke in Type Two Diabetic Rats Improves Vascular Permeability Measured by MRI. PLoS One. 2016;11:e0149147. doi: 10.1371/journal.pone.0149147

39. Cui C, Ye X, Chopp M, Venkat P, Zacharek A, Yan T, Ning R, Yu P, Cui G, Chen J. miR-145 Regulates Diabetes-Bone Marrow Stromal Cell-Induced Neurorestorative Effects in Diabetes Stroke Rats. Stem Cells Transl Med. 2016;5:1656–1667. doi: 10.5966/sctm.2015-0349

40. Venkat P, Zacharek A, Landschoot-Ward J, Wang F, Culmone L, Chen Z, Chopp M, Chen J. Exosomes derived from bone marrow mesenchymal stem cells harvested from type two diabetes rats promotes neurorestorative effects after stroke in type two diabetes rats. Exp Neurol. 2020;334:113456. doi: 10.1016/j.expneurol.2020.113456

41. Di Meglio L, Solo Nomenjanahary M, Bedoucha L, Dupont S, Zemali F, Ollivier V, Journe C, Jandrot-Perrus M, Rambaud T, Mazighi M, et al. Neutrophil extracellular traps-targeting therapy with deoxyribonuclease 1 reduces large vessel occlusion-induced downstream microvascular thromboinflammation in a rat model of stroke. Research and Practice in Thrombosis and Haemostasis. 2025;9:103206. doi: 10.1016/j.rpth.2025.103206

42. Di G, Vázquez-Reyes S, Díaz B, Peña-Martinez C, García-Culebras A, Cuartero MI, Moraga A, Pradillo JM, Esposito E, Lo EH, et al. Daytime DNase-I Administration Protects Mice From Ischemic Stroke Without Inducing Bleeding or tPA-Induced Hemorrhagic Transformation, Even With Aspirin Pretreatment. Stroke. 2025;56:527–532. doi: doi:10.1161/STROKEAHA.124.049961

43. Mouloud Y, Staubach S, Stambouli O, Mokhtari S, Kutzner TJ, Zwanziger D, Hemeda H, Giebel B. Calcium chloride declotted human platelet lysate promotes the expansion of mesenchymal stromal cells and allows manufacturing of immunomodulatory active extracellular vesicle products. Cytotherapy. 2024;26:988–998. doi: 10.1016/j.jcyt.2024.04.069

44. Dominici M, Le Blanc K, Mueller I, Slaper-Cortenbach I, Marini FC, Krause DS, Deans RJ, Keating A, Prockop DJ, Horwitz EM. Minimal criteria for defining multipotent mesenchymal stromal cells. The International Society for Cellular Therapy position statement. Cytotherapy. 2006;8:315–317. doi: 10.1080/14653240600855905

